# DiatomNet: An automatic Diatom genus identification system through microscopic images and Deep Learning

**DOI:** 10.1101/2025.02.10.635050

**Authors:** Pedro Villar, Jacobo Casado, David Fernández, Pedro Sánchez, Siham Tabik

**Affiliations:** Department of Software Engineering, University of Granada, Granada, 18071, Andalucia, Spain; Department of Computer Science and Artificial Intelligence, University of Granada, Granada, 18071, Andalucia, Spain; Andalusian Research Institute in Data Science and Computational Intelligence (DASCI), University of Granada, Granada, 18071, Andalucia, Spain; Department of Botany, Campus Fuentenueva. University of Granada, Granada, 18071, Andalucia, Spain

**Keywords:** Diatoms classification, Microscope images, Convolutional neural networks, Deep Learning

## Abstract

Diatoms are microscopic organisms belonging to the algae kingdom. They adapt to the ecosystem and modify their shape and texture depending on hundreds of ecosystem variables. Hence, these micro-organism are considered as the most accurate indicator to measure water quality. Commonly, the recognition of the class of diatoms in a microscopic image has always been done by expert biologists knowledgeable about the morphometric characteristics of these organisms. This work proposes a new automatic diatom genus recognition system from microscopic images using state-of-the-art deep CNNs. In particular, 1) we developed a public high quality database organized into 44 genus-level diatom classes, 2) designed a robust diatom classification model, and 3) provided a user-friendly interface to utilize our automatic diatom recognition tool.

## 1. Introduction

Diatoms are microscopic organiscvvvffms belonging to the algae kingdom. These unicellular organisms form the most common group of microorganisms in aquatic habitats and their presence in the food chain of aquatic ecosystems is vital. Diatoms adapt to the ecosystem and modify their shape and texture depending on hundreds of ecosystem variables.

They have several advantages over others biondicators: they have a quick response to the ecosystem conditions variation and they are very sensitive to changes in the environment that other bioindocators do not detect, especially in many of the chemical variables of water, such as pH, where diatoms are capable of measuring changes in the order of thousandths of a millisecond (1). All these characteristics make them the most accurate indicator to measure water quality.

They are also easy to sample without making a big impact in the ecosystem which make them ideal for complex studies related with climatic change and its evolution (2; 3). Commonly, the recognition of the diatoms genus in a microscopic image has always been done by expert biologists knowledgeable about the morphometric characteristics of these organisms, mainly length, wide and shape, and internal details of the diatom, like striae density (texture information) (see an example in Figure 1). Once the features are identified, the classification is performed manually by comparing with diatoms previously classified.

**Figure 1:**
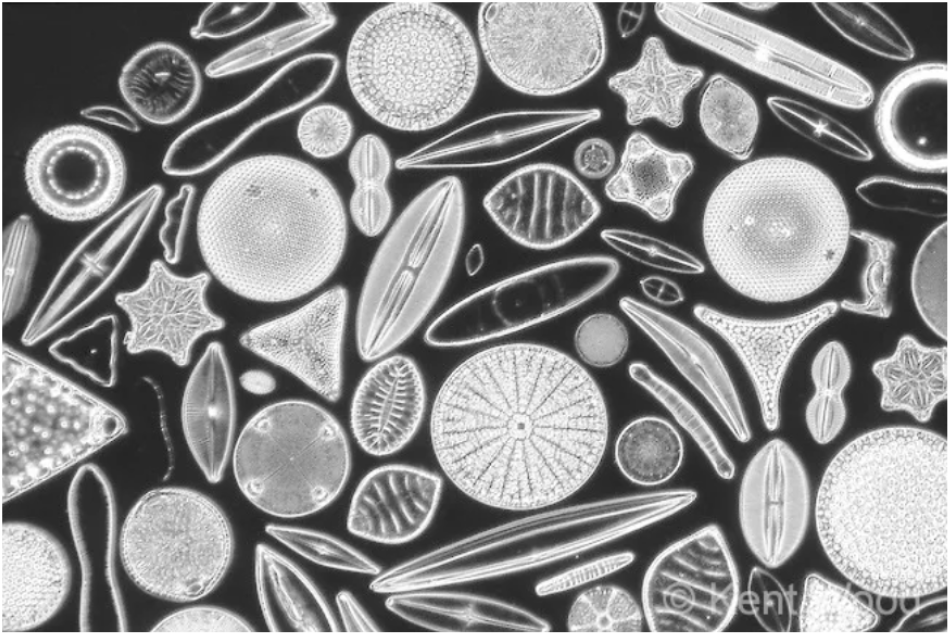
Several diatoms genus in a microscope sample. Source: (7).

Several studies intended to ease diatom genus recognition but usually using features extracted manually (4) or considering a very small number of diatom classes (5). To the best of our knowledge, there does no exist any *end-to-end* automatic system for recognizing diatom genus from images.

Thanks to the last advances in deep learning models and the creation of new massive databases, such as ImageNet^1^, deep Convolutional Neural Networks (CNNs) have shown to be robust for the extraction of spatial patterns from images (6). Actually, CNNs continue to be one of the most appropriate network architectures in all fundamental computer vision tasks, e.g., image classification and object detection in images.

The full automatization of diatom genus classification from microscopic image is challenging for the next reasons:

- Expert biologists are required to determine new key features analysis when a new genus appears.
- Manual analysis of diatoms images is a long and tedious process due to the elevated number of features to analyze. In addition, for water quality studies at least 400 images have to be analyzed (8).
- There does not exist any database for training automatic diatom genus classification model.
- Diatoms have inter-genus similarities and intra-genus dissimilarities that make the classification task more difficult. Figure 2 shows two diatoms very similar visually but belonging to two different genera. Whereas, Figure 3 shows three diatoms with visually different shapes but belong to the same genus.
- According to (9), there are currently around 10.000 recognized species, however the estimated total number of species is greater than 200.000.
- Diatoms changes shape during their life cycle. This polymorphism make the manual classification very difficult (10).

**Figure 2:**
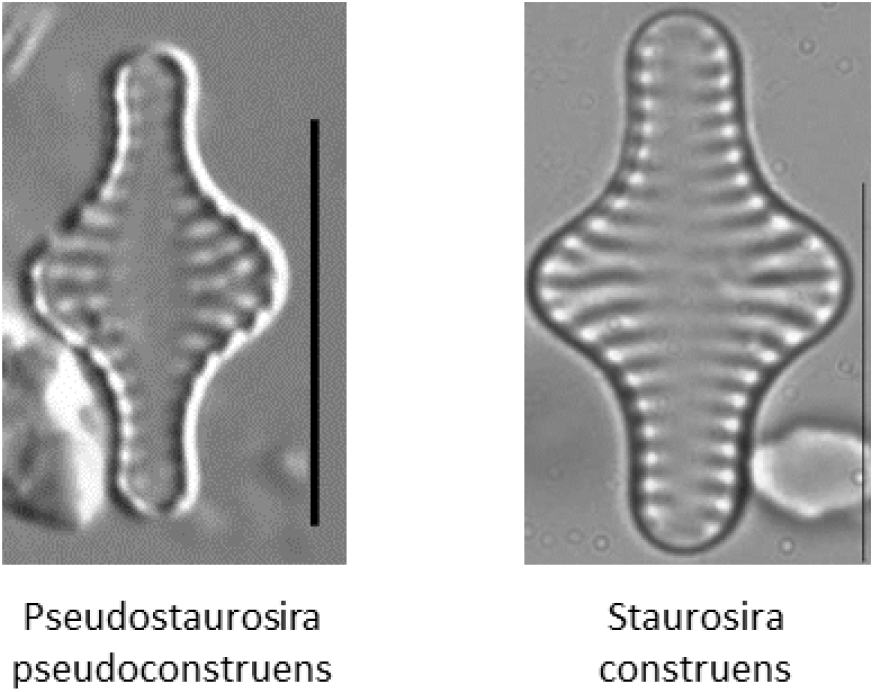
Two diatoms of different genera. Source: (11)

**Figure 3:**
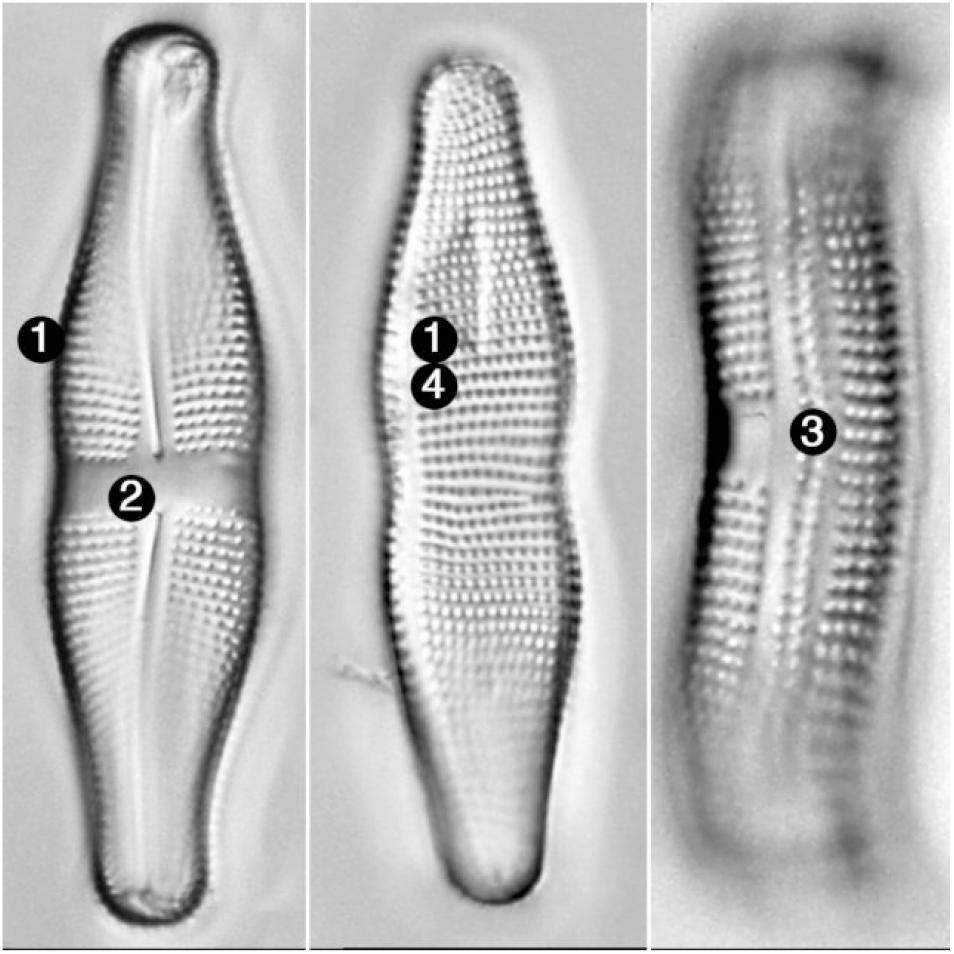
Images of diatoms from genus *Achnantes*. Source: (11).

This work focuses on developing an automatic diatom class recognition system from microscopic images using state-of-the-art deep CNNs. Specifically, 1) we developed DiatomNet, a public smart database of microscopic images organized into 44 genus-level diatom classes, 2) we built a robust model for identifying the diatom genus that includes a data preprocessing strategy to improve the robustness and generalization capacity of the classification model, and 3) we provided a user-friendly interface that utilizes our model, creating an automatic diatom recognition tool.

The rest of this paper is organized as follows. Section 2 presents an overview on CNNs, including the architectures and techniques used in this work. In section 3 a summary of previous related works for classifying diatoms from images is given. In section 4.1 we present the new dataset, Diatom- Net, created in this work. Our automatic classification model of diatoms based on microscopic images is presented in section 5. Section 6 describes the automatic tool developed for diatom recognition. Finally, conclusions are stated in Section 7.

## 2. Convolutional Neural Networks for image classification

CNNs are a widely used type of artificial neural network. They constitute the state-of-the-art in object recognition and classification in images. CNNs are able to learn a large number of features from an image through a number of hidden layers. They apply several transformations to the data using its principal operation named convolution. In this operation, an input block is transformed into an output block by means of a kernel. Each convolutional layer has different number of kernels. The input of a convolutional layer is convoluted with each kernel. The values of these kernels, called weights, are automatically learned during the training process. At the end of the learning process, these values encode the distinctive features of the class.

To increase the non-linearity of the models, every convolutional layer is followed by a non-linear operation, typically the Rectified Linear Unit (ReLU) operation. To increase the abstraction level of the obtained features, a pooling layer is used to reduce the size of the feature maps obtained from the convolutional layer. The pooling layers allow the network to extract from simple in low levels to complex features to higher levels of the network.

In 2019, a new architecture called EfficientNet was proposed (12). The main purpose of EfficientNet is scaling the CNNs using a dynamic strategy to obtain better results and efficiency. In 2021, the authors proposed a new version, EfficientNetV2 that incorporates several improvements both in performance and calculation speed.

Usually, CNNs require large datasets to obtain good results. To overcome this constraint several improvement techniques are applied, like *transfer learning* and *data augmentation*.

*Transfer learning* consists of, instead of training the network from scratch, retraining a network previously trained on a different problem, preferably related to the current problem. Depending on the size of the training dataset, one can retrain only the last layers of the network, the part that converts the high level features into the corresponding class. This process is known as *fine tuning*.

*Data augmentation* consists of increasing the size of the training set artificially by applying several transformations to the original images like flipping them horizontally or vertically, rotating them, shifting them, etc. In practice, the user can choose how much distortion to apply (zoom 50% of the image, rotate the image 45 degrees, etc.). It is necessary to be careful with this technique as the images may lose their meaning (13).

One of the limitations of CNNs is their poor interpretability capacity. The weights are adjusted to minimize the error function, but the model does not explains what criteria the network has taken into account to decide that a particular image belongs to a class, since CNNs have hundreds of thousands of interconnected neurons that come together to make a final decision. Improving the interpretability of neural networks is a current field of research. There exist several tools, such as *Saliency Map* (14), LIME (15) o Grad- CAM++ (16), that try to explain the predictions of a network in a more interpretable way for the human.

## 3. Related work on diatoms classification

This section presents a brief review of previous works on classification of diatom images, using from manual classification to semiautomatic and finally, automatic classification of diatoms.

### 3.1. Manual classification of diatoms

Ten years ago, the manual classification of diatoms was the only way to classify diatom images. The usual procedure consisted of analyzing each diatom individually by extracting its key features (generally, morphometric characteristics), and comparing with the key features of each class. In the experiments performed in (17), it was verified that the ability of an expert to identify correctly a diatom is about 80%. This technique is quite sensitive to microscope light as some details may not be visible, requiring the application of post-processing techniques to visualize all parts of the diatom. Another drawback of this procedure is the complexity of adding new classes of diatoms, since experts are needed to describe the key features of the new class.

### 3.2. Semiautomatic classification of diatoms

Due to the complexity of the manual task, semiautomatic classification methods have been developed. Information about these methods can be found in (18) and (19). Usual techniques such as *holografic filters* (20) and *harmonic decomposition* (21) are highlighted. These semiautomatic methods aid to automate the process but require human intervention. We briefly describe two methods of this category:

The first methods was proposed in (4). The authors used an image processing software called **ImageJ** to compare a target diatom image with a database composed of reference images of one diatom for each class of interest. The system gets two characteristic of the diatom, the outline and the ornamentation (information about the interior patterns of the diatom). This task requires human intervention. Once the pattern and ornamentation have been obtained, these two values are automatically compared with the values of the database images and the diatom with the most similar image in the database is the output of the method.

**ADIAC** (*Automatic Diatom Identification and Classification*) (22) was the first attempt to apply machine learning algorithms to identify and classify diatoms. In this work, a great and well labeled database were built and diatoms morphometric descriptors were identified. These descriptors were used as features in machine learning models. They managed to collect 4, 700 images of 500 species of diatoms and they achieved an accuracy of 96.7% using *random forest*. It was necessary to use 321 mathematical descriptors of the diatoms to differentiate the classes. The process is not completely automatic as human intervention is required to add new species of diatoms.

### 3.3. Automatic classification of diatoms

The aim herein is to fully automate the diatoms classification process.

Very few works addressed diatoms classification using Deep Learning. The authors in (5) consider a limited number of genera (20) and performs the classification at the species level, at which the diatoms characteristic are more dissimilar. A recent study (23) used Mask R-CNN and U-Net models to perform an automatic binary pixel-wise segmentation of diatoms in diatom microscopy slides in form of gigapixel-sized virtual slide images. The proposal of this work is not identifying diatoms but instead classifying a pixel into either a diatom pixel or background. This kind of methods can be combined to ours as a pre-processing phase, so that first a diatom is located in an input tile then identified using our proposed classification model.

In the present work, we consider more than 40 diatoms genera and more than 900 species. Our method classifies at the genus level, which is challenging, as it has to overcome the inter-genus similarities and intra-genus dissimilarities commented in Section 1.

## 4. Methodology

In this paper, we first built a new database called Diatom- Net especially suitable for training Deep Learning models and analyzed several EfficientNetV2 based architectures to build a diatoms classifier. We also compare with AlexNet (24), one of the most popular CNNs. To improve the generalization capacity and robustness of the model we used data-augmentation for balancing the classes. We also analyzed the impact of transfer learning on the generalization capacity of the model and used LIME tool for inspecting the quality of the predictions.

### 4.1. DiatomNet database: a new diatom microscopic images dataset

A large number of studies have proven that building higher quality models requires higher quality data, also known as Smart Data (25). The concept of Smart Data refers to the process of converting raw data into higher quality data with higher concentration of useful information (26). Smart data includes all pre-processing methods that improve value and veracity of data. Examples of these methods include noise elimination, data-augmentation (27) and data transformation (28) among other techniques.

In the context of microscopic images the bias can come from different sources. Each laboratory has a different microscope model which affects the resolution, intensity and quality of the final image. To avoid this bias, the used images have been collected from different laboratories.

In this section we describe the construction of the database, a preliminary data analysis and the pre-processing techniques used. Our DiatomNet is provided through this link: (29)

### 4.2. Construction of DiatomNet database

In a first phase, we have collected images of 44 genera of diatoms to solve a classification problem of 44 classes. However, it is important to emphasize that the model can be readjusted to consider a larger number of classes at any time. Moreover, if new images from an existing class are obtained, it is possible to retrain the model to further improve its prediction quality.

The set of diatoms genera selected to build Diatom- Net together with the number of images per each genus showed are in Table 1. These images have been obtained from the following sources: Diatoms.org (11), ANSP (30), AQUALITAS (31) and the images provided by the ADIAC project, found on RGBE website (32).

**Table 1.**
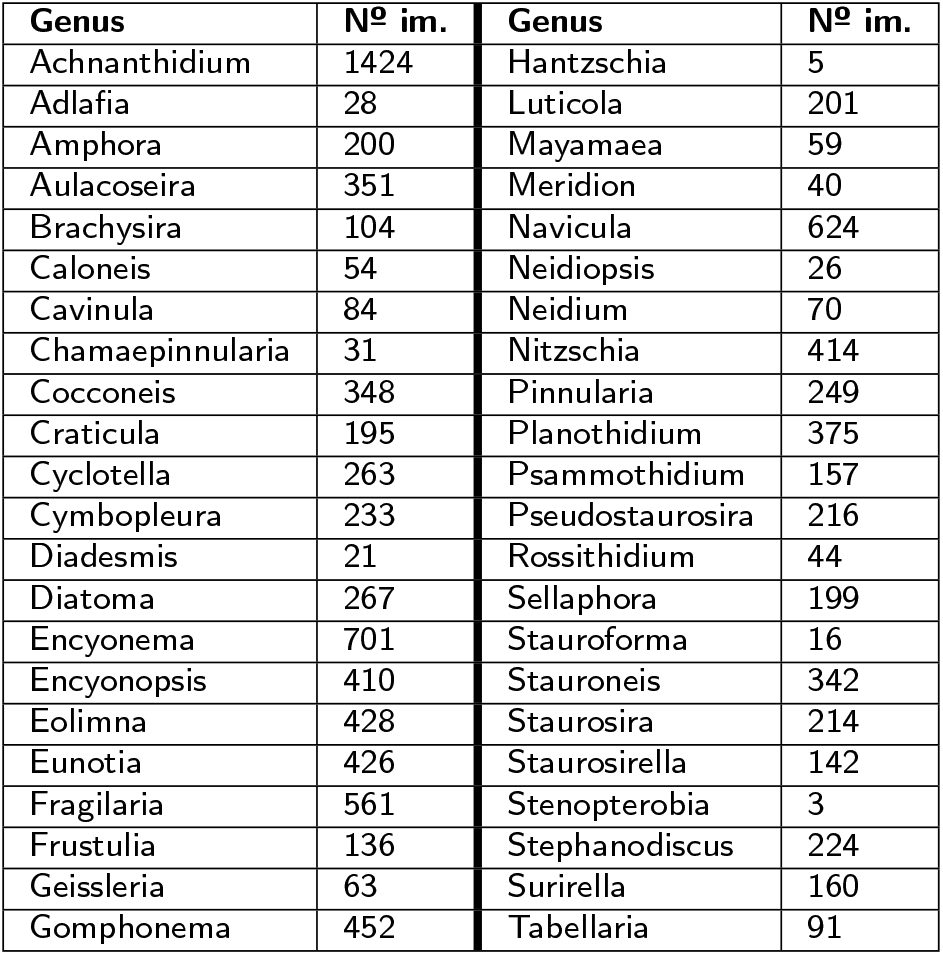
Number of images (№ im.) of each genus of the 44 classes of DiatomNet database.

### 4.3. Preliminar analysis of DiatomNet

After the construction process, we obtained a dataset of 10, 650 images organized into 44 classes. The origin and quantity of the dataset are shown in Figure 4 and Table 1.

**Figure 4:**
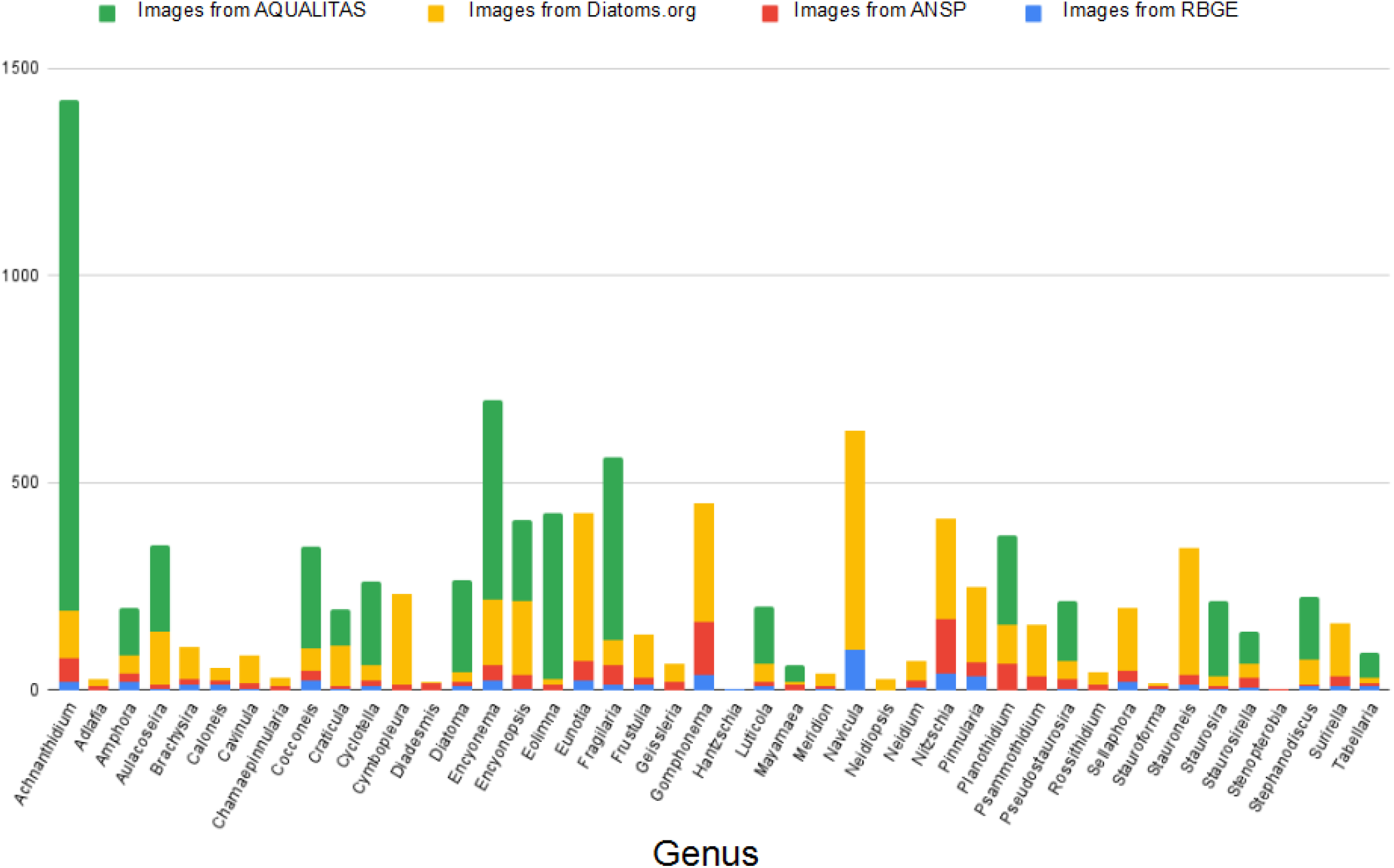
Number of images obtained according to class and colored according to origin (graphic legend on top).

As can be seen in Table 1 and Figure 4, most classes are represented by reasonable number of samples except two, class *Hantzschia* is represented by 3 and class *Stenopterobia* by 5. As it is not practical to train a model with so few data, these two classes are eliminated from this study, but they could be added in the future if new images are obtained. Therefore, we finally used 10, 642 images. Most of the images have been obtained from AQUALITAS and Diatoms.org. The obtained dataset is unbalanced, some classes are represented by 20 images while others include 600. This problem will be treated at the end of this section, in the preprocessing techniques subsection.

The included images are highly heterogeneous as they were captured by different microscopes in different environments and hence they have different zoom levels, size of the diatom within the image, light intensity, etc. See examples in Figure 5.

**Figure 5:**
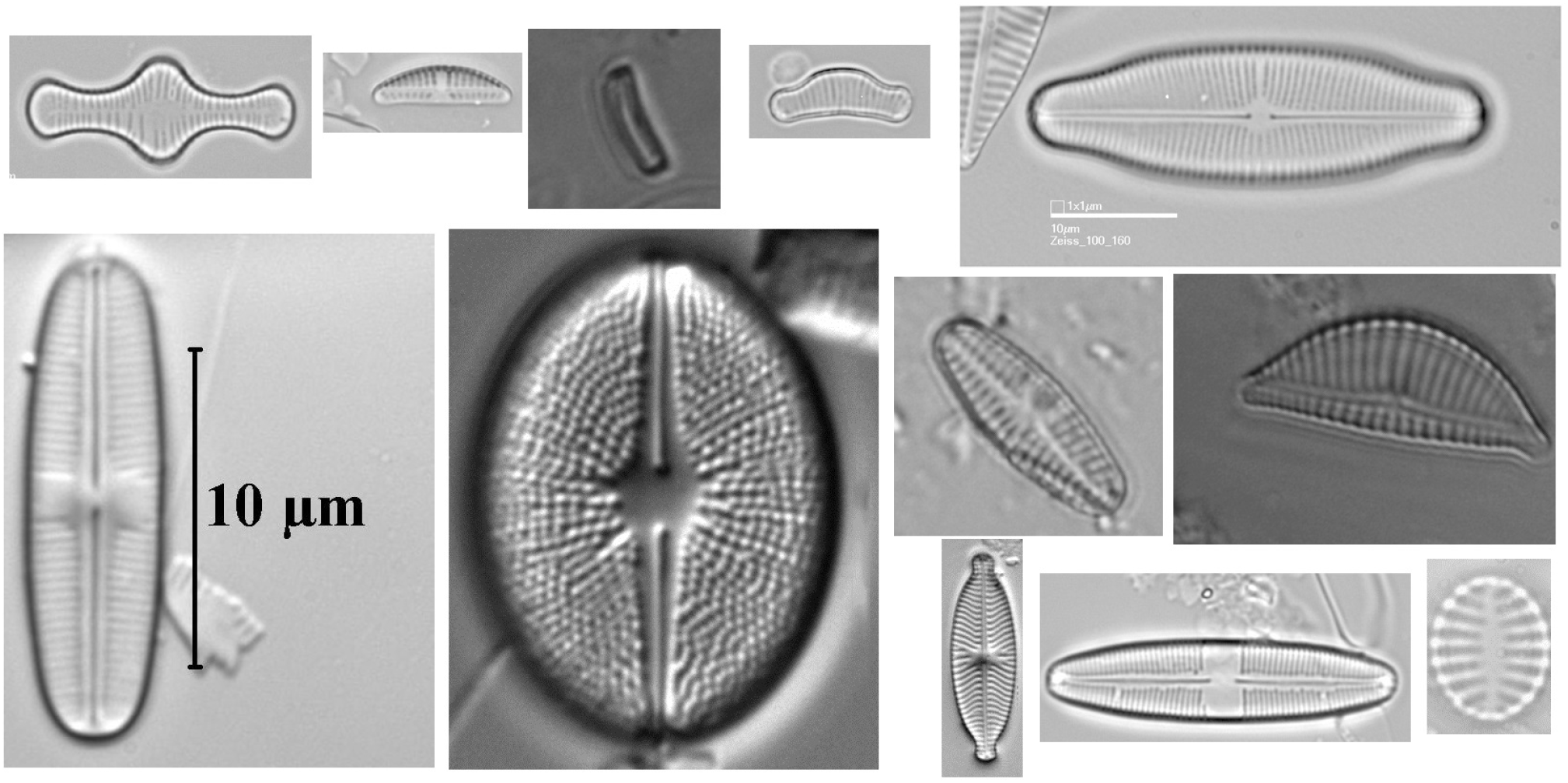
A sample of 12 diatom images selected randomly from the built dataset.

As it can be seen, it is necessary to eliminate the scale information produced by the electron microscope software in most samples. Such noise can cause the network to try to associate the shape of that information as a relevant feature and lose generalization capacity with images that do not have these marks. On the contrary, the pieces of dead diatoms or pieces of dust in the background can be interesting for improving the generalization capacity of the model. If models are trained with images that contain a totally clean background, the network could consider any piece of dust or dirt as a diatom and worsen the prediction capacity.

Furthermore, the differences in diatoms size can be clearly observed. There are images in which the diatom completely covers the image and others where the diatom occupies less than 30% of the image. It is beneficial for the learning process that diatoms occupy at least 60% of the image, so the images that are small will be rescaled so that they meet that minimum.

It can be seen that the images do not have the same light intensity. Besides, in most images diatoms are displayed vertically and horizontally although some of them are rotated. Analyzing the datasets, it can be seen that all diatoms from the images of RBGE, Diatoms.org and ANSP are displayed vertically or horizontally while the images of AQUALITAS include rotated diatoms with different angles. Overall, in the majority of the images, the diatoms are displayed vertically or horizontally.

### 4.4. Preprocessing techniques

In this section We describe all the preprocessing techniques used in this work: the cleaning and trimming process, the use of data augmentation for improving the behavior of the model and the use of data augmentation for balancing the classes.

#### 4.4.1. Cleaning and cropping process

The cleaning and cropping process is a manual process, image by image, in which it is detected if the image needs to be cropped and if it has any marks generated by the microscope around the diatom. This process could not be automated because each image has a different size and the mark is generated depending on the location of the diatom in the image and the technique used by the laboratory. An image editor software has been used to remove the marks, using an intelligent fill tool that takes into account the areas around the mark to replace it with similar pixels, simulating being part of the original image. An example of this preprocessing is shown in Figure 6, where after removing the microscope marks from the image, part of it has been cut to adapt it to the diatom, so that it occupies most of the image.

**Figure 6:**
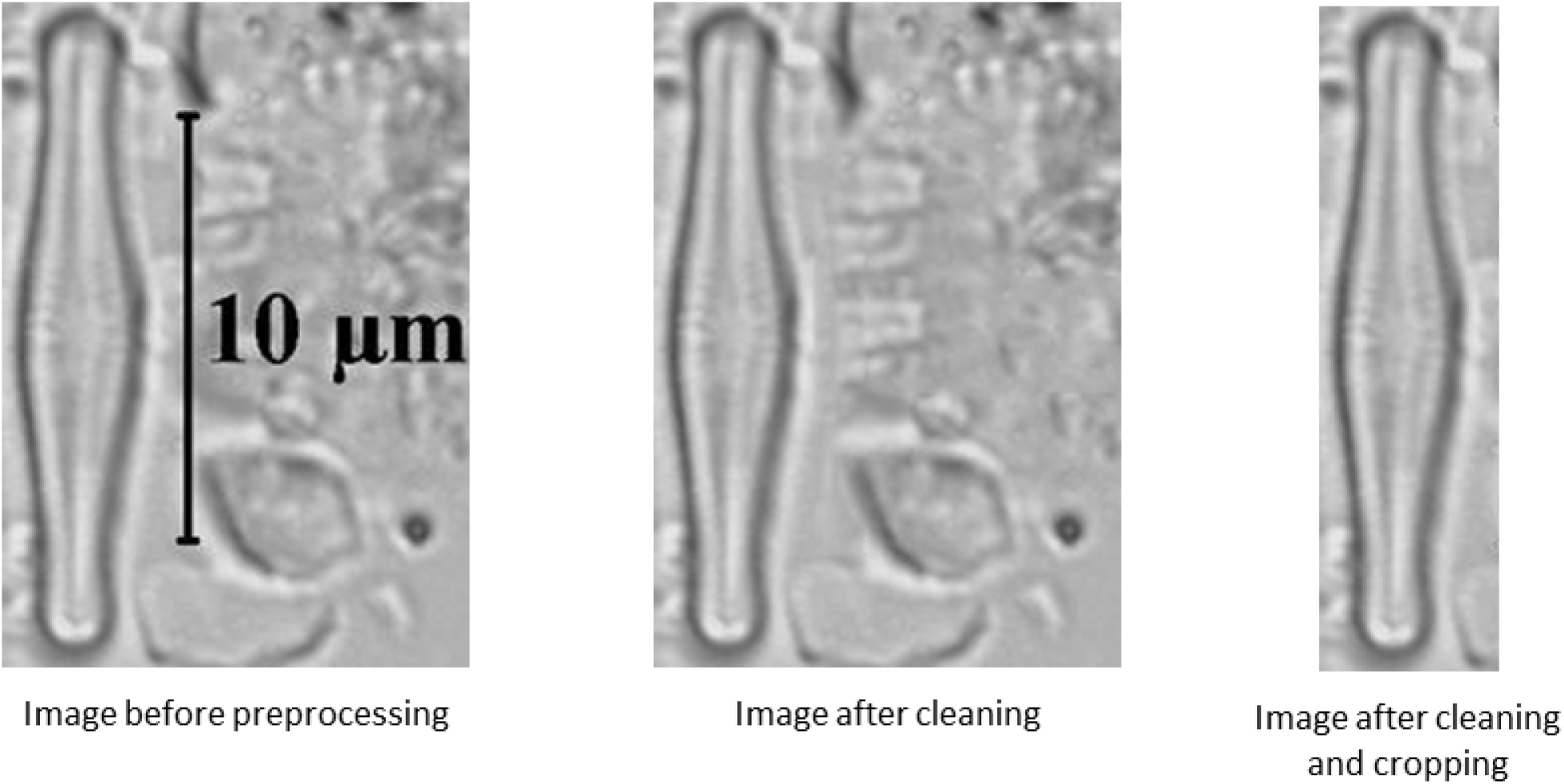
Image preprocessing example

In the dataset, there are images that require one of the two steps and others that require both or none depending the their quality. The marks of dirt and remains of diatoms have been maintained in the background of the images, as this context information can be valuable for training the network.

#### 4.4.2. Improve generalization and robustness via data augmentation

As commented previously, the lack of data is the main problem of CNNs. The main idea of data augmentation is to help with this problem by increasing the quantity and also the variety of data through artificial or synthetic images generated from the original data. Such artificially generated data can be seen as data extracted from a very close distribution to the original distribution, thus obtaining new data that is not exactly the same as the original, but very similar.

If new artificial images are generated from an original image by applying basic transformations such as rotations or zoom, the network will extract more general characteristics from that data, since it must be able to classify that object regardless of the angle and lighting of the original image, creating some invariance to all these situations that can occur in the context of an image. In this work, this technique helps to reduce the bias generated in each laboratory, which uses different techniques and microscopes to take the images.

In addition, it has been shown that performing data augmentation helps avoiding overfitting, creating models that generalize better on new data, as well as being invariant to changes in the image (33). Figure 7 shows an image of the dataset to which basic data augmentation has been applied: a random rotation in the range of 0 to 180 degrees, a change in image intensity in the range of 70 to 130%, and a random vertical or horizontal mirroring.

**Figure 7:**
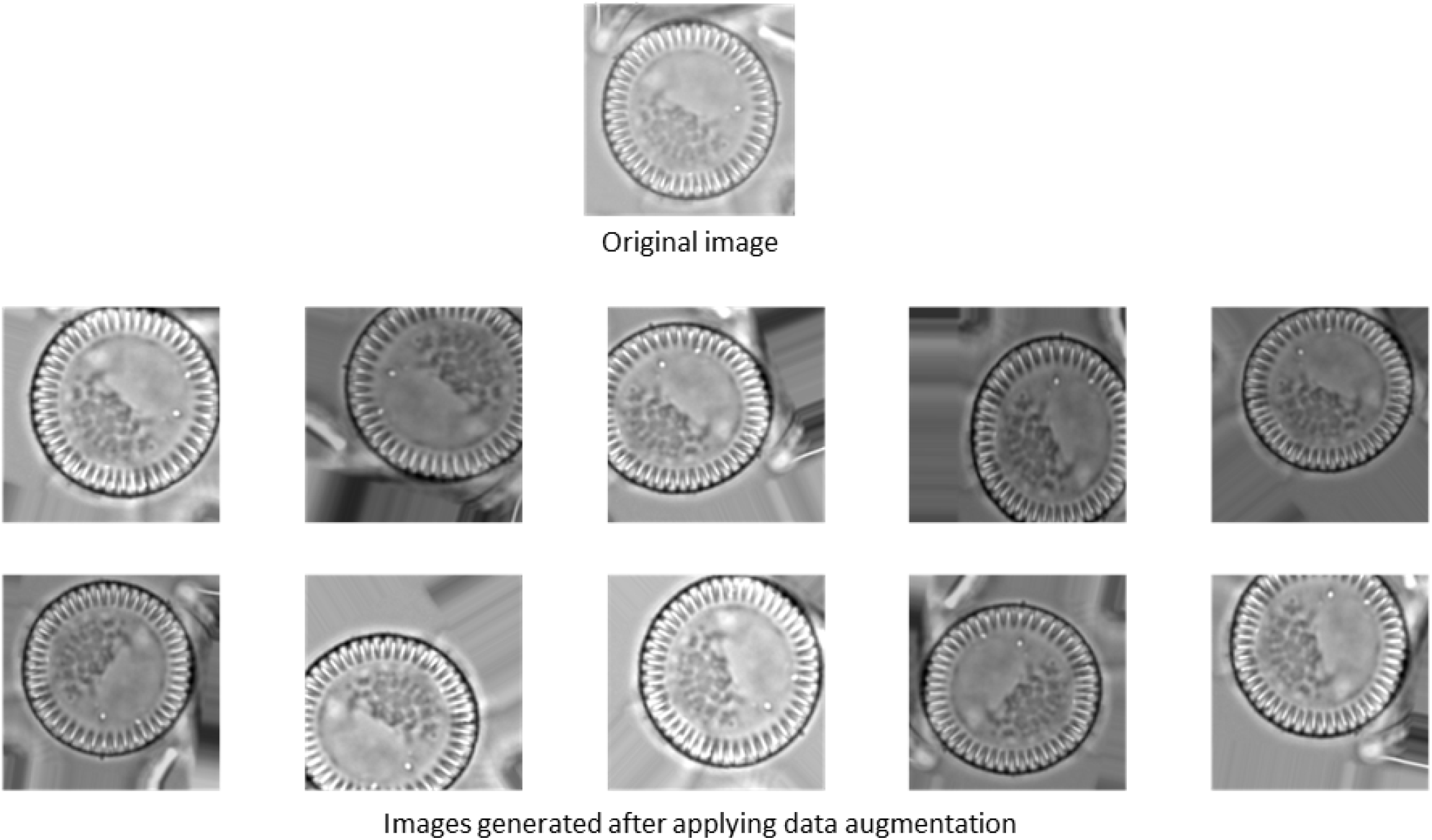
Artificially generated images from the original image using data augmentation

#### 4.4.3. Balancing data using data augmentation

The high unbalance degree of DiatomNet was commented in Section 4.3. Some minority classes have on the order of 20 to 30 images, while majority classes have 200 to 600 images. This situation generates a bias towards the major classes, causing the network to capture the characteristics of the classes with more images, leaving the classes with fewer images marginalized. To avoid this problem, we propose to balance the minority classes through data augmentation, making all classes have the same number of images in the training process.

## 5. Automatic classification of diatoms using microscopic images

We propose the use of CNNs for the automatic classification of diatoms by means of microscopic images. Next, we describe the experimental framework, the experimental results, a detailed analysis of the best model, a brief study on the explainability of that model and the learning of the final model used for the deploy in the automatic recognition tool built.

### 5.1. Experimental framework

To build a robust diatom classifier, we evaluated several EfficientNetV2 based architectures (34). Figure 8 shows the architecture of EfficientNetV0, the baseline for the family of EfficientNetV2 networks. For comparison purpose, we also used AlexNet (24).

**Figure 8:**
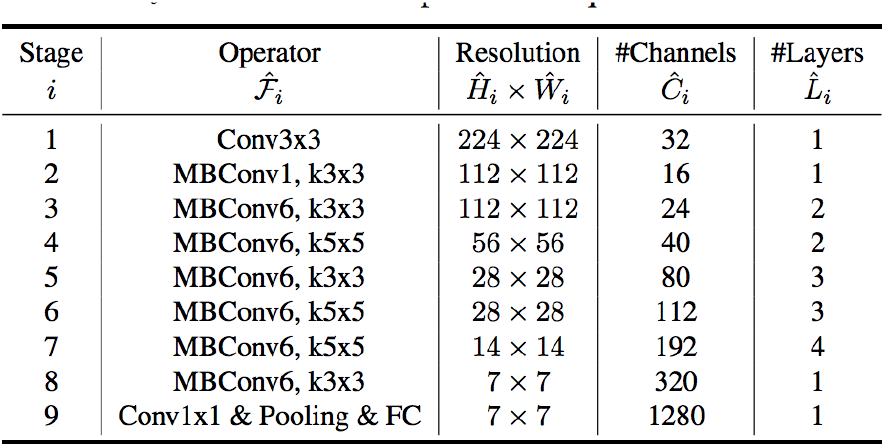
Architecture of EfficientNetV0 (12).

We have used Google colab running Python notebooks (vs. 3.6.9) for the datasets creation and models training. The main software libraries used are:

- NumPy (vs. 1.15.4)
- Pandas (vs. 0.24.2)
- TensorFlow (vs. 2.8.2)
- Keras (vs. 2.2.4)
- Scikit-Learn (vs. 0.21.2)
- OpenCV (vs. 4.0.0)

The accuracy metric is employed for the experimental evaluation. The dataset is divided in training set (80%) and test set (20%), maintaining the same distribution of examples for each class in both sets (argument *stratified* in the function used for dividing the dataset). The size of those two datasets are **8174** and **2468** images respectively. We applied a 3-fold cross validation in the learning process. So, the training set is divided in three sets with the same size and we perform three experiments using one fold as validation set and the other two folds joined as usual training data set.

We have used an input image size of 224 × 244 pixels, a fairly common format in most models trained on ImageNet. All images have been resized to this size. Table 2 shows the hyperparameters used.

**Table 2.** Set of hyperparameters.

|  | Num. epochs | Batch size | Learning rate |
| --- | --- | --- | --- |
| EfficientNet | 100 | 16 , 32 | 0.001 |
| AlexNet | 100 | 16 , 32 | 0.001 |

Regarding the class balancing, data augmentation was used and we have performed experiments with two values of the number of images for all the classes: 400 images and 1000 images.

Callbacks are extra options that can be applied to the model to control its behavior during the training phase. They help to get better results and simplify the training task. In this work, we have used three callbacks:

- **EarlyStopping**: This callback allows to detect when there is overfitting by monitoring the validation set and ending the training phase when the model gets worse a number of times in a row. This number of consecutive times without improvement is the parameter called *patience*, which we have set to **6**. Another parameter of this callback called *restore_best_weights* has been set to **True** to retrieve the model weights before the worsening.
- **ModelCheckpoint**: It saves a copy of the best model found considering the metric stored in *monitor*. In order to improve the generalization capability, we have chosen the best model on the validation set, so that *monitor* = **“val_loss”**
- **ReduceLROnPlateau**: This callback monitors a metric in each epoch (in our case, again over the validation set). If the metric does not improve in a number of epochs set by the parameter *patience*, the hyperparameter “learning rate” is reduced by a proportion set in the parameter *factor*. In the experiments, we use *patience* = **2** and *factor* = **0.5**.

### 5.2. Experimental results

Table 3 shows the obtained mean accuracy in training and validation for some representative models. By analyzing this table, one can observe that AlexNet achieves lower accuracy although it includes more parameters than EfficientNet models. This can be explained by the fact that AlexNet is a primitive architecture that does not include several modern optimizations.

**Table 3.** Results obtained for the cross validation in each architecture. It shows the mean of accuracy in training and validation for the three folds

| Architecture | Train | number of parameters | Balancing | $acc_{mean}$ train | $acc_{mean}$ val |
| --- | --- | --- | --- | --- | --- |
| AlexNet | *From scratch* | 46.924.586 | 400 | 0.77643 | 0.72479 |
| EfficientNetB0 | *Transfer learning* | 53.802 | 400 | 0.92929 | 0.85558 |
| EfficientNetB0 | *From scratch* | 4.061.350 | 400 | 0.99470 | 0.94334 |
| EfficientNetV2B0 | *From scratch* | 5.912.506 | 400 | 0.99589 | 0.94476 |
| EfficientNetV2B0 | *From scratch* | 5.912.506 | 1000 | 0.99876 | **0.97202** |
| EfficientNetV2B0 | *From scratch* | 5.912.506 | NO | 0.98749 | 0.94490 |
| EfficientNetV2B1 | *From scratch* | 6.913.854 | 1000 | 0.99893 | 0.94293 |
| EfficientNetV2S | *From scratch* | 20.231.290 | 1000 | 0.99963 | 0.92293 |

On the other hand, the family of EfficientNetV2 networks show good behavior, adjusting almost completely to the training data and presenting no overfitting (the difference between the mean accuracy in training and validation is very small). These CNNs perform better when trained from scratch and when no transfer learning is applied. This means that the generic ImageNet feature extractor are not enough to extract the features of such a specific domain. It can be seen that the largest networks of this family (EfficientNetV2B1 and EfficientNetV2S) start to show signs of a little overfitting. Considering the EfficientNetB0 networks, the results of version 2 (“V2B0”) are slightly better than those of the previous version (“B0”).

The model with the best generalization capability (best mean accuracy in validation set) was trained with a balanced dataset with 1000 images per class. Notice that the results from training the same model without class balancing or balanced at 400 images per class are very close. From these results one can also observe that a model trained with 10 to 30 images of minority classes behaves similarly to a model trained with 400 examples per class. This can be explained by the fact that the samples of majority classes in the validation sets are distributed vertically or horizontally, as commented in Section 4.3 and hence the characteristics learned by the models with augmented data are not included in this validation set, only vertical and horizontal feature extractors are being used. In real situations this will not be the case and there will be surely diatoms rotated at any angle. It would be interesting to repeat the validation process without retraining the models but rotating each image of the validation sets in a random range from 0 to 180 degrees, obtaining a data set more faithful to reality. In this way, it can be verified whether a real improvement has actually been obtained in the models trained with more images.

Table 4 shows the results with the new validation set. All models have behaved a little worse, but the models that have decreased the most have been those trained without data augmentation, and therefore without class balancing or those trained with low rate of data augmentation. The images which the model has been validated are exactly the same ones as before but rotated slightly. The great impact on the improvement of the model can be seen when training with artificially created images to make the network robust to this type of transformations. This robustness obtained by the model due to the use of artificial data can be extrapolated to other types of transformations, such as the luminosity or the zoom of the image.

**Table 4.**
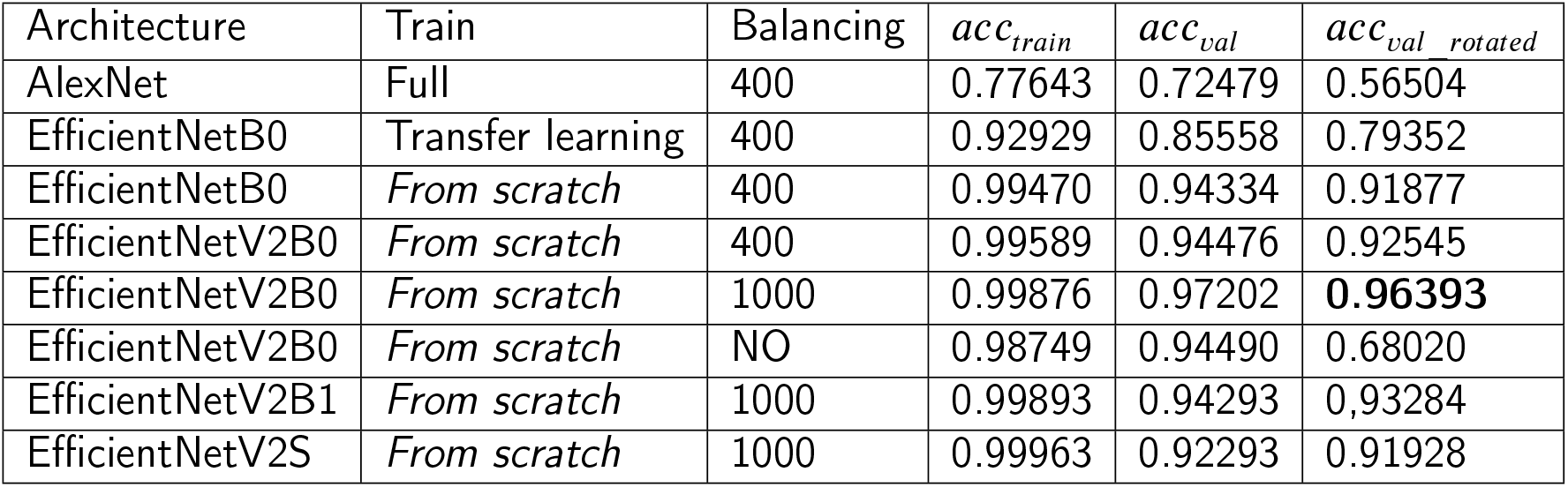
Results obtained in each model after performing cross validation with the validation set randomly rotated.

### 5.3. Analysis of the winning model

The winning model of the previous comparison is EfficientNetV2B0 model obtained by trainning the full network on 1000 images per class. We choose the model trained with folds 1 and 2 (validated with fold 0) because it is the model with the best results (accuracy of 97.018% in fold 0).

We now use the test set (section 5.1) to validate the model with images that were not seen during the construction process of the model. The accuracy over the test set is 94.89%. If the test set is randomly rotated as in the previous training phase, the accuracy is 94.21%. To check the performance of the model at the class level we analyzed the confusion matrix and the classification report for the test set.

Figure 9 shows the confusion matrix. It can be seen that there is not a great confusion between classes. The **Achnanthidium** class is the one with the largest number of confusions, but it is also the one with the largest number of images. The two classes that are most confused with each other are **Achnanthidium** and **Eolimna**.

**Figure 9:**
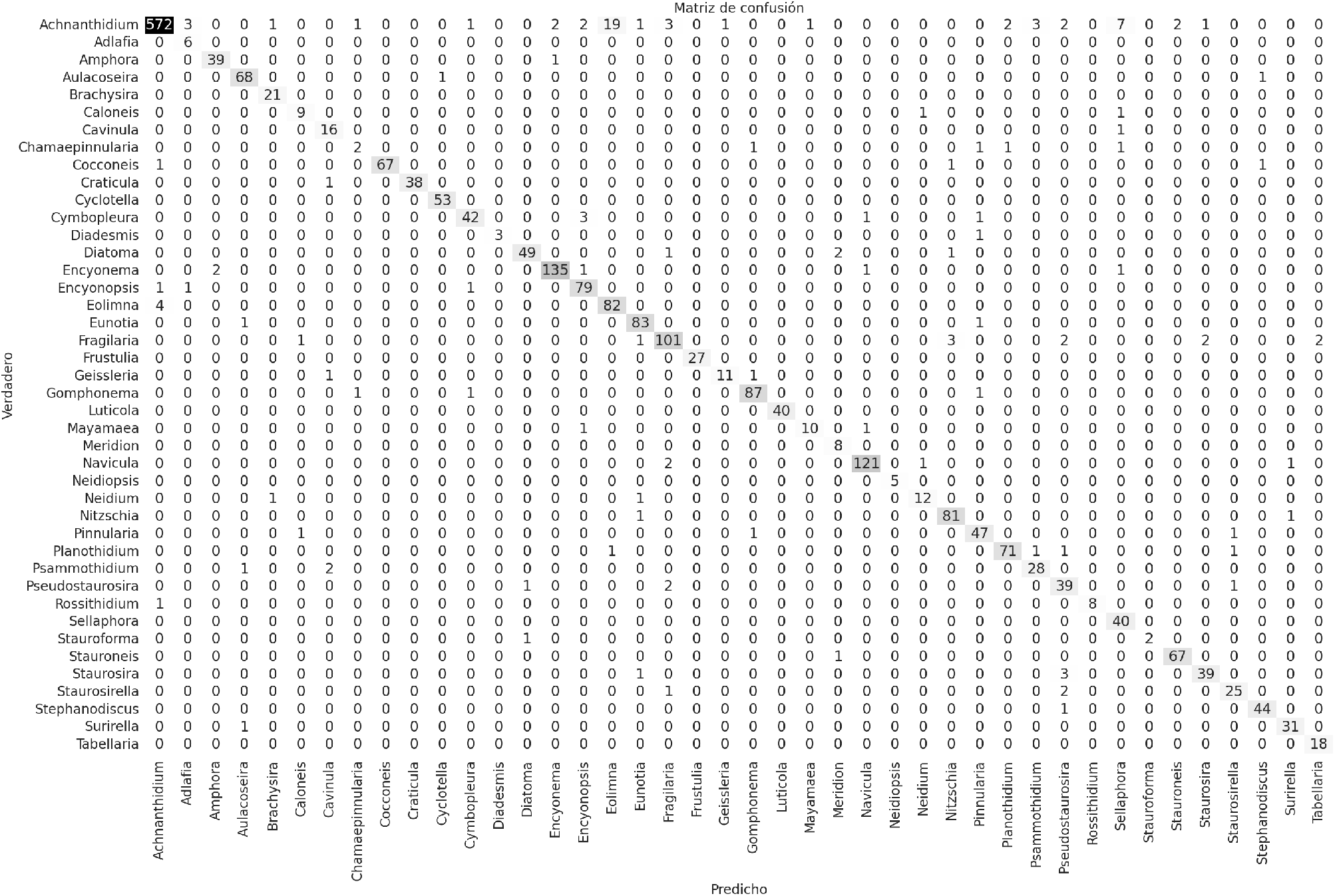
Confusion matrix of the test set.

An example of this confusion is shown in Figure 10. The images in this case are low quality. It is possible that the model is not able to find differentiating characteristics between the classes as the diatoms in the image are not well defined. To avoid such confusion it is recommended to collect and use only high-quality images without noise and blurriness.

**Figure 10:**
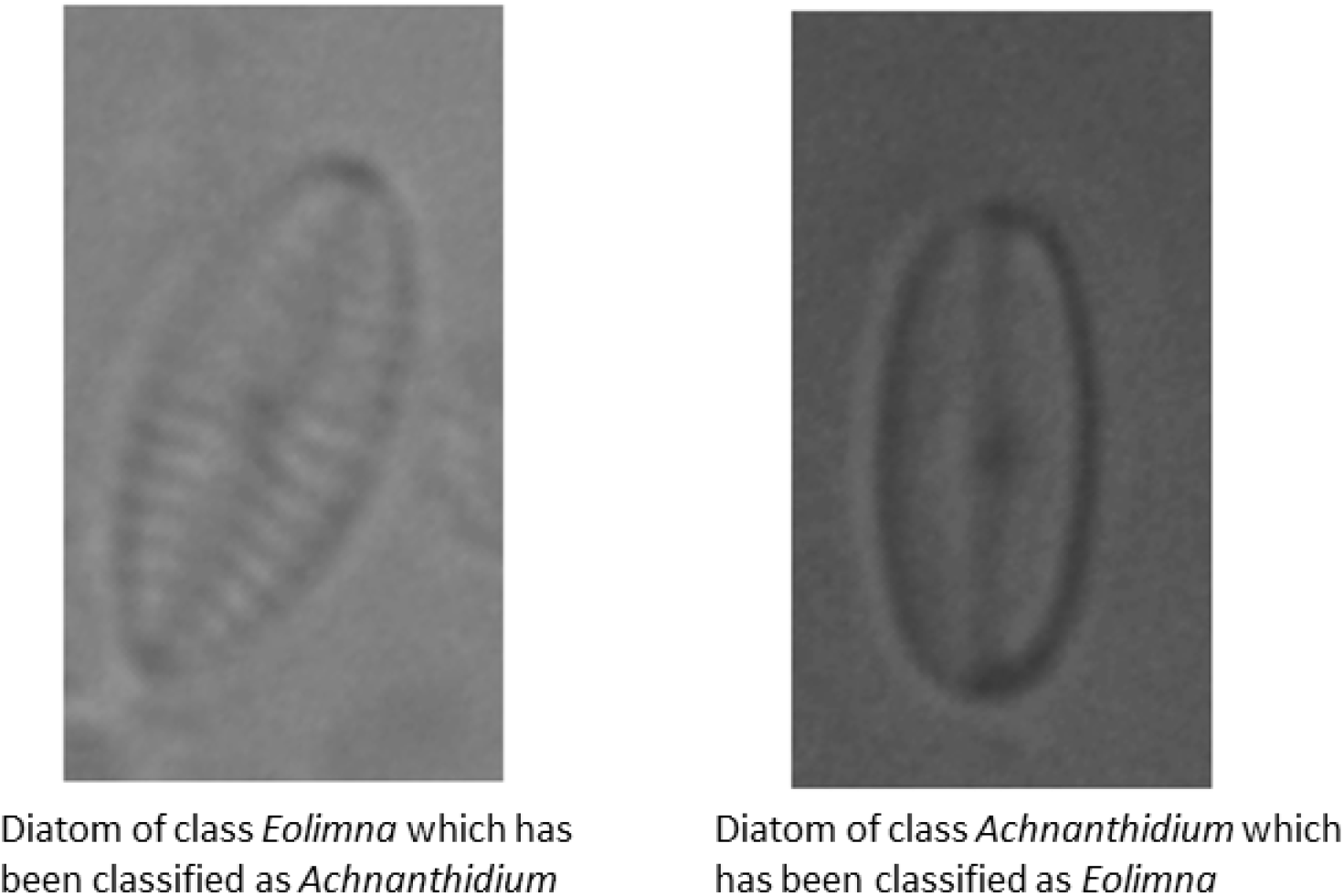
Example of two misclassified images. Source: (31)

The classification report of the winning model is showed in Table 5. It can be seen that the mean of *recall* and *precision* have values above 90%. In addition, most of the classes have a behavior close to the average in both metrics, which indicates that the behavior of the model at the class level is quite uniform. The Stauroforma and Neidiopsis classes show perfect behavior at the class level with a *precision* of 100% and a *recall* of the same value despite being ones of the least represented in the dataset. Other classes such as Luticola, Craticola or Neidium replicate this behavior with a greater number of images. The model has been able to extract representative characteristics of a large number of classes, which have allowed them to be completely differentiated from the rest.

**Table 5.**
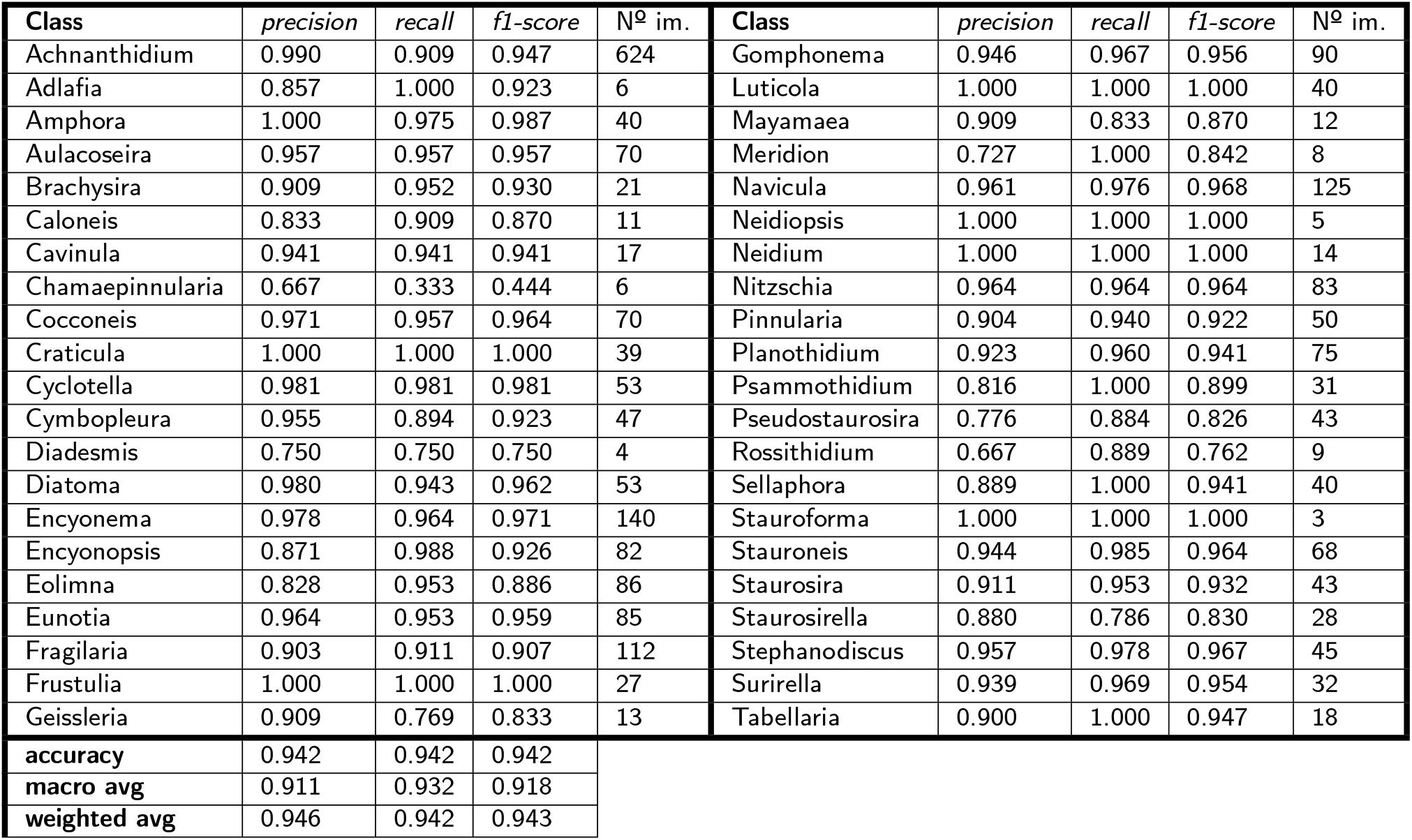
classification report for the test set of the winning model.

| Class | precision | recall | f1-score | Nº im. | Class | precision | recall | f1-score | Nº im. |
| --- | --- | --- | --- | --- | --- | --- | --- | --- | --- |
| Achnanthidium | 0.990 | 0.909 | 0.947 | 624 | Gomphonema | 0.946 | 0.967 | 0.956 | 90 |
| Adlafia | 0.857 | 1.000 | 0.923 | 6 | Luticola | 1.000 | 1.000 | 1.000 | 40 |
| Amphora | 1.000 | 0.975 | 0.987 | 40 | Mayamaea | 0.909 | 0.833 | 0.870 | 12 |
| Aulacoseira | 0.957 | 0.957 | 0.957 | 70 | Meridion | 0.727 | 1.000 | 0.842 | 8 |
| Brachysira | 0.909 | 0.952 | 0.930 | 21 | Navicula | 0.961 | 0.976 | 0.968 | 125 |
| Caloneis | 0.833 | 0.909 | 0.870 | 11 | Neidiopsis | 1.000 | 1.000 | 1.000 | 5 |
| Cavinula | 0.941 | 0.941 | 0.941 | 17 | Neidium | 1.000 | 1.000 | 1.000 | 14 |
| Chamaepinnularia | 0.667 | 0.333 | 0.444 | 6 | Nitzschia | 0.964 | 0.964 | 0.964 | 83 |
| Cocconeis | 0.971 | 0.957 | 0.964 | 70 | Pinnularia | 0.904 | 0.940 | 0.922 | 50 |
| Craticula | 1.000 | 1.000 | 1.000 | 39 | Planothidium | 0.923 | 0.960 | 0.941 | 75 |
| Cyclotella | 0.981 | 0.981 | 0.981 | 53 | Psammothidium | 0.816 | 1.000 | 0.899 | 31 |
| Cymbopleura | 0.955 | 0.894 | 0.923 | 47 | Pseudostaurosira | 0.776 | 0.884 | 0.826 | 43 |
| Diadesmis | 0.750 | 0.750 | 0.750 | 4 | Rossithidium | 0.667 | 0.889 | 0.762 | 9 |
| Diatoma | 0.980 | 0.943 | 0.962 | 53 | Sellaphora | 0.889 | 1.000 | 0.941 | 40 |
| Encyonema | 0.978 | 0.964 | 0.971 | 140 | Stauroforma | 1.000 | 1.000 | 1.000 | 3 |
| Encyonopsis | 0.871 | 0.988 | 0.926 | 82 | Stauroneis | 0.944 | 0.985 | 0.964 | 68 |
| Eolimna | 0.828 | 0.953 | 0.886 | 86 | Staurosira | 0.911 | 0.953 | 0.932 | 43 |
| Eunotia | 0.964 | 0.953 | 0.959 | 85 | Staurosirella | 0.880 | 0.786 | 0.830 | 28 |
| Fragilaria | 0.903 | 0.911 | 0.907 | 112 | Stephanodiscus | 0.957 | 0.978 | 0.967 | 45 |
| Frustulia | 1.000 | 1.000 | 1.000 | 27 | Surirella | 0.939 | 0.969 | 0.954 | 32 |
| Geissleria | 0.909 | 0.769 | 0.833 | 13 | Tabellaria | 0.900 | 1.000 | 0.947 | 18 |
| accuracy | 0.942 | 0.942 | 0.942 |  |  |  |  |  |  |
| macro avg | 0.911 | 0.932 | 0.918 |  |  |  |  |  |  |
| weighted avg | 0.946 | 0.942 | 0.943 |  |  |  |  |  |  |

### 5.4. Explainability of the model using LIME

In this section the LIME tool (15) is applied to see what the model has focused on when making its decisions. The tool has been used in cases in which the model correctly classifies the image (true positives) and cases in which the model fails, to assess its predictions in both situations.

Figure 11 displays the explanations provided by LIME for two examples of model correct predictions. It can be seen that in both predictions, the model looks at the internal structure of the diatom and extracts very high-level features (in this case, corresponding to the ornamentation) to make its decision. In this way, it has managed to correctly classify the images.

**Figure 11:**
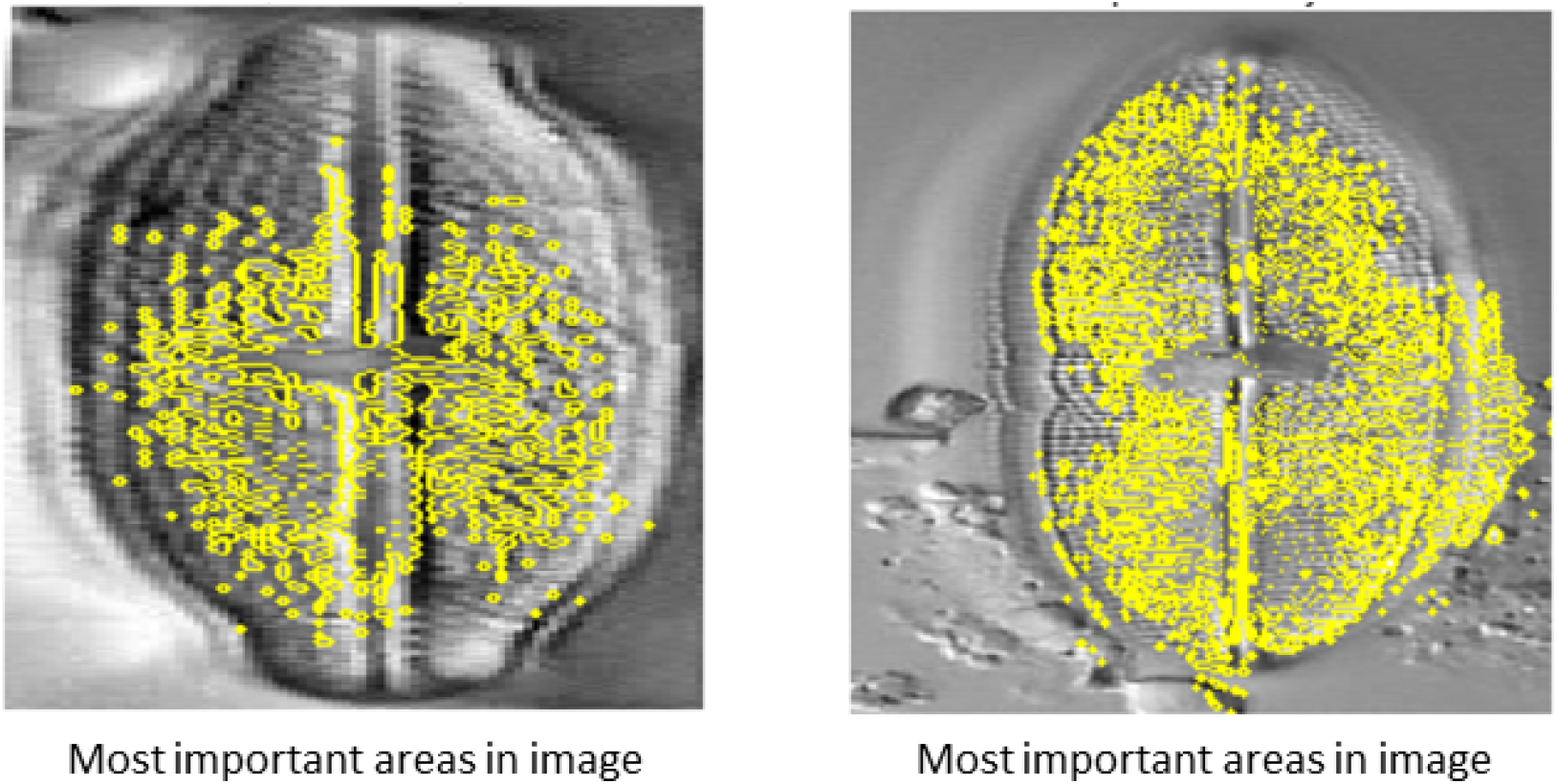
Explanations provided by LIME for correctly classified images

Figure 12 shows two examples of images misclassified by the model together with its explanation. In both cases, the model has not been able to capture the distinctive characteristics of the diatom, and therefore ends up misclassifying it.

**Figure 12:**
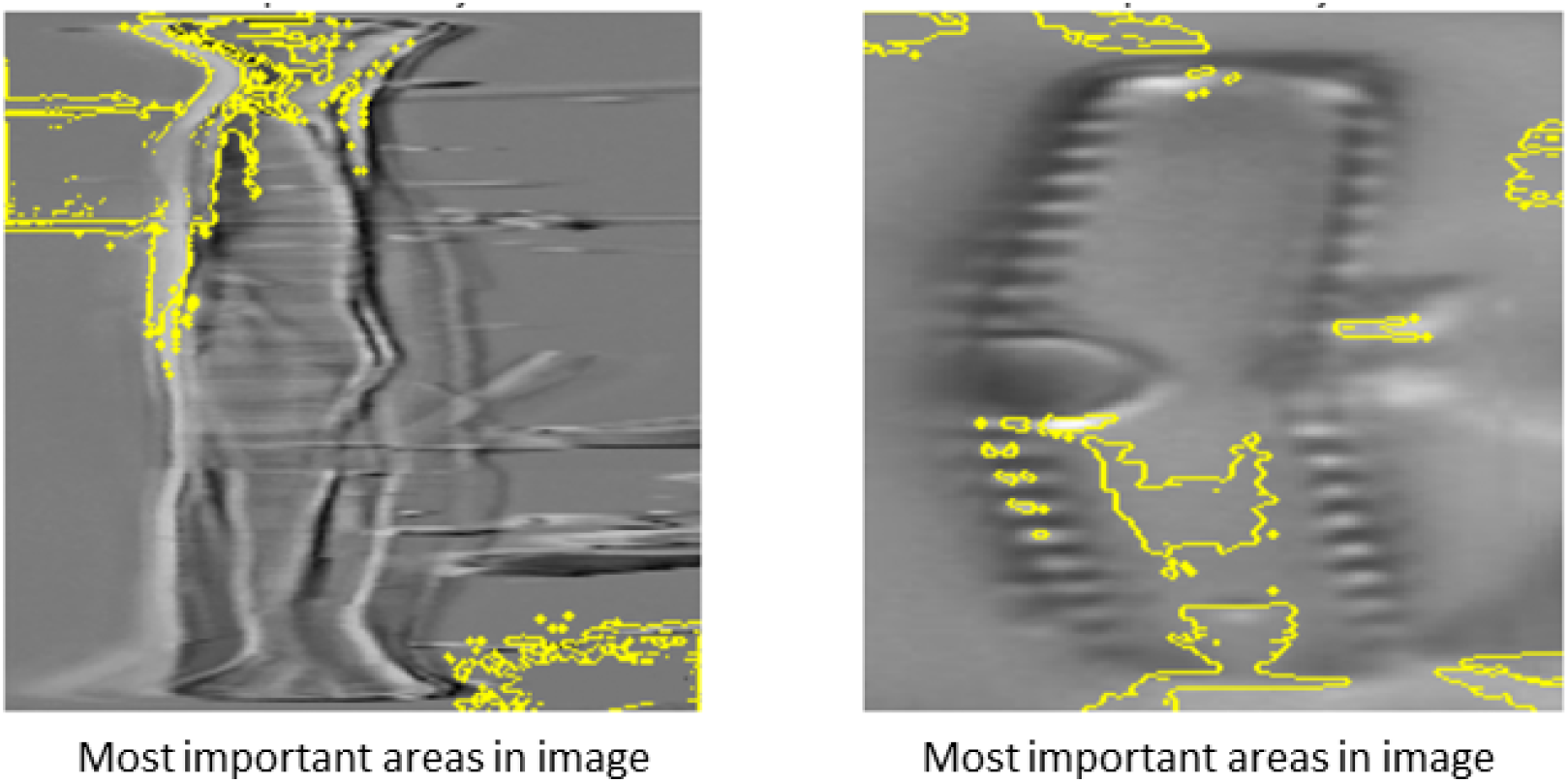
Explanations provided by LIME for misclassified images

Given the great utility of the LIME tool, it allows inspecting and evaluating the decisions of the used models in a more visual way.

### 5.5. Getting the final model for deployment with an interface

Finally, we retrain the model using all the database (including the testset). The obtained model has been implemented in a graphical interface as an automatic classification tool. The validation set is still essential to avoid overfitting and to stop the training. Therefore, a 20% of the total dataset is still reserved for the validation set.

The best model achieves an accuracy of 95.48% in the validation set. Analyzing the evolution of the model throughout the training, it can be seen that the overfitting is very slight, which can be further reduced by retraining the model on new images.

## 6. Automatic diatoms recognition tool

The final model has been implemented in a simple web page, obtaining a graphical interface deployed in a local web. With this tool, the user provides a diatom image and get a prediction about the genus (class) of the diatom. In this section, we describe the process of the interface creation and show an example.

### 6.1. Installation

A *python* file called **app.py** is used, where a local server is running using **Flask** in the address **http://127.0.0.1:5000**. All the information necessary for the installation can be found at (35).

The idea of this web page is to serve as a simple and reproducible tool so that anyone can run it quickly, so some files have been prepared so that the tool can be portable to any device. There exists the possibility of customizing the appearance of the web page so that it is more intuitive for the user.

All the files are in a repository (35), where the user manual is also attached.

### 6.2. Example of using the tool

After accessing the web page, the web interface is showed in Figure 13.

**Figure 13:**
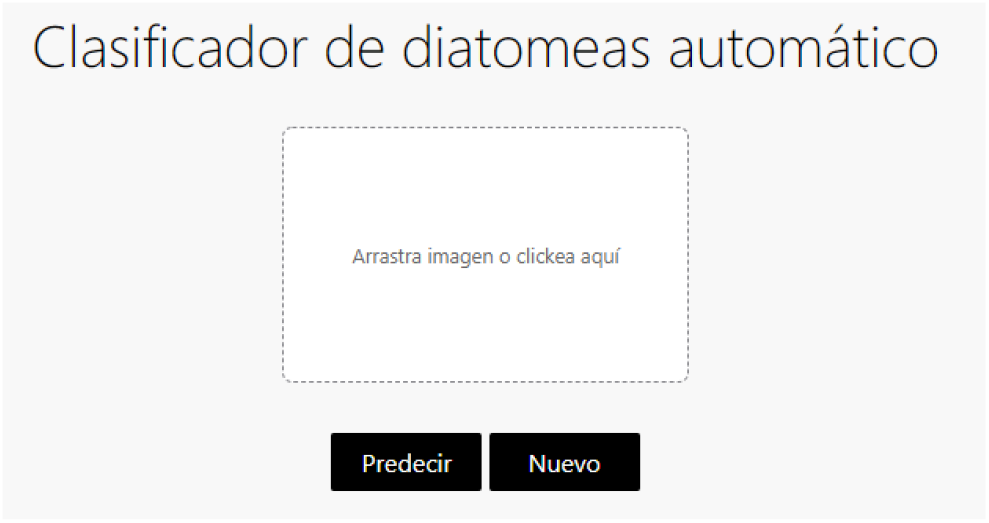
Web page interface made for interaction with the end user

The user can click on the square or drag to provide the image of the interest to the automatic classification system. After pressing the predict button, the system returns the prediction as shown in the Figure 14. In this way, the user can consult the five classes with the greatest similarity with the input image.

**Figure 14:**
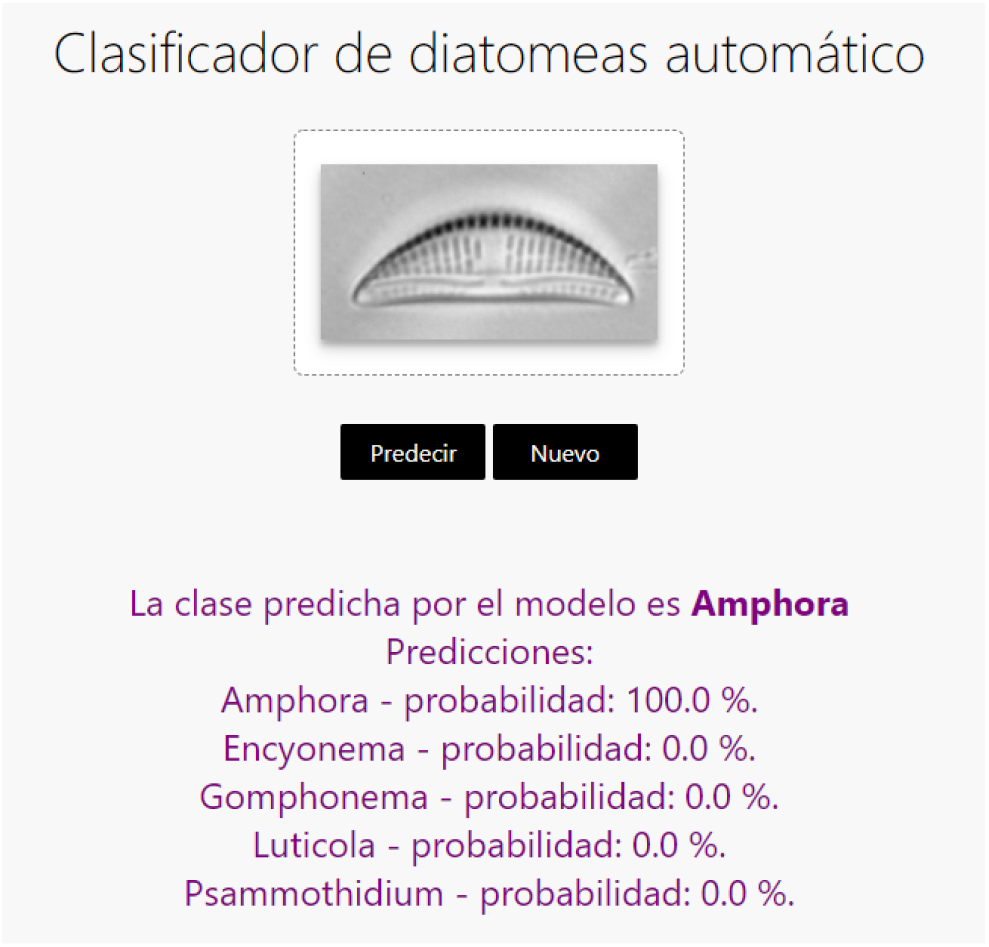
Example of prediction obtained by the tool

## 7. Conclusions

In this paper a new method for automatic diatoms genus identification based on microscopic images using CNNs is proposed. We built a new dataset with diatoms microscopic images (DiatomNet database), obtained from various sources. We have applied several preprocessing techniques to improve the generalization capacity and robustness of the model. Various versions of the state-of-the-art CNN architectures have been analyzed, obtaining a model with good prediction ability as it correctly classifies 95,48% of the test images. This model is expected to be improved in the future using more images coming from various sources. We finally present a web graphical interface that predicts the genus of a diatom based on its image.

## Acknowledgement

This research is part of the project “Thematic Center on Mountain Ecosystem & Remote sensing, Deep learning-AI e-Services University of Granada-Sierra Nevada” (LifeWatch-2019-10-UGR-01), which has been co-funded by the Ministry of Science and Innovation through the FEDER funds from the Spanish Pluriregional Operational Program 2014-2020 (POPE), LifeWatch-ERIC action line.

